# Comparative single-cell lineage bias in human and murine hematopoietic stem cells

**DOI:** 10.1101/2024.11.19.624262

**Authors:** Isaac Shamie, Meghan Bliss-Moreau, Jamie Casey Lee, Ronald Mathieu, Harold M. Hoffman, Bob Geng, Nathan E. Lewis, Yanfang Peipei Zhu, Ben A. Croker

**Author notes:** Correspondence: Ben A. Croker,; Yanfang Peipei Zhu,; Nathan E. Lewis. Contributed equally (Isaac Shamie, Meghan Bliss-Moreau, Jamie Casey Lee). Contributed equally (Nathan E. Lewis, Yanfang Peipei Zhu, Ben A. Croker). The authors declare no financial conflicts of interest.

## Abstract

The commitment of hematopoietic stem cells (HSC) to myeloid, erythroid, and lymphoid lineages is influenced by microenvironmental cues, and governed by cell-intrinsic and epigenetic characteristics that are unique to the HSC population. To investigate the nature of lineage commitment bias in human HSC, mitochondrial single cell (sc) ATAC-Sequencing (mt-scATAC-Seq) was used to identify somatic mutations in mitochondrial DNA to act as natural genetic barcodes for tracking the *ex vivo* differentiation potential of HSC to mature cells. Clonal lineages of human CD34^+^ cells and their mature progeny were normally distributed across the hematopoietic lineage tree without evidence of significant skewing. To investigate commitment bias *in vivo*, mice were transplanted with limited numbers of long-term HSC (LT-HSC). Variation in the ratio of myeloid and lymphoid cells between donors, although suggestive of a skewed output, was not altered by increasing numbers of LT-HSC. These data suggest that the variation in myeloid and lymphoid engraftment is a stochastic process influenced by the irradiated recipient niche and not a cell-intrinsic lineage bias of LT-HSC.

## Introduction

Hematopoietic stem cells (HSC) are classically considered to have the capacity for complete regeneration of the hematopoietic compartment. More recent analyses indicate additional complexity and heterogeneity in the HSC compartment, with lineage-restricted or lineage-biased HSC considered a feature of mammalian hematopoiesis.^1–13^ Central to the concept of lineage bias is an assumption that cells used for studying HSC commitment are HSC and not multipotent progenitors or lineage-committed progenitors. Changes in differentiation of cells downstream of the LT-HSC must also be evaluated when considering the potential lineage bias of a LT-HSC. Functional validation of these heterogeneous characteristics minimally requires demonstration of the capacity to transplant single HSC to lethally-irradiated or hemoablated recipients.^14^ Retrospective analysis of single HSC cells injected into recipient mice and their progeny can validate the isolation procedure, and support data obtained from protocols for prospective isolation of these rare bone marrow cell populations. Next generation sequencing (NGS) and gene expression analyses have identified differentially expressed genes modulated specifically within the LT-HSC population, and enabled development of antibodies for purification and phenotypic analysis.^15–18^ However, successful engraftment of mice with single HSC still remains challenging. Many studies aiming to repopulate mice with HSC report variable reconstitution using inconsistent definitions of engraftment including both survival and >0.1-1% repopulation in any lineage, suggesting that some HSC isolation protocols result in the purification of lineage-restricted progenitor cells and mature hematopoietic cells.

HSC tagging methodologies also provide a means to study the *in vivo* function of HSC. Clonal relationships between hematopoietic stem progenitor cells (HSPC) and mature hematopoietic lineages have been explored using inducible DNA ‘‘barcoding’’ methodologies during embryogenesis or postnatal life, genetically modified mice expressing fluorescent markers within the HSC population or upon the transplantation of virally-transduced barcoded single HSCs to lethally-irradiated or hemoablated adult murine or non-human primate recipients.^19–25^ These methods have provided a wealth of data supporting the role of hematopoietic progenitor cells in driving steady state hematopoiesis and the apparent lineage bias of LT-HSC, but also strong counterarguments.^26,27^ HSPC differentiation can also be tracked by mapping somatic mutations in the mitochondria, which act as natural barcodes for cells.^28^ This modified scATAC-seq method (mt-scATAC-seq) captures both open nuclear chromatin and mitochondrial DNA sequence to identify somatic variants.

In this study, mt-scATAC-seq was used to exploit the somatic mutations in mitochondrial DNA as natural barcodes and thus track the *ex vivo* differentiation of human CD34+ HSC. Furthermore, mt-scATAC-seq was used to generate a data set allowing the comparison of human CD34+ cells lineage differentiation *in vitro* with mouse LT-HSC differentiation *in vivo*. In this, we found that clonal lineages of human CD34^+^ cells and their mature progeny are normally distributed across the hematopoietic lineage, and no evidence of significant skewing of downstream cell types was observed. When single LT-HSC were transplanted into mice, we found limited variation in the ratio of myeloid and lymphoid cells between donors. Therefore, variation in myeloid and lymphoid engraftment is a stochastic process influenced by the irradiated recipient niche and not cell-intrinsic differences in LT-HSC.

## Results

### mt-scATAC-seq defines clonal lineages in human CD34+ cells

To study differentiation of primary human CD34^+^ cells to committed myeloid and erythroid cells, CD34^+^ cells from two healthy donors were cultured for 72h in the presence of recombinant human SCF/IL-3/IL-6/Flt3L/G-CSF/GM-CSF. Donor CD34^+^ cells were multiplexed for single cell capture and library preparation before (input) or after culture (Figure 1A, Table S1, see Methods). Cells from an additional six healthy donors were analyzed to increase the power to delineate lineage relationships before and after cell culture. Using scATAC-seq, mitochondrial DNA somatic variants were identified and further used to generate clone-specific “natural” barcodes. These mitochondrial barcodes are inherited in clonal hierarchies to define clonality of CD34^+^ stem cells and their progeny. In parallel, the lower-coverage reads in nuclear open-chromatin regions were used for assigning cell lineages by assessing differential peaks in gene regulatory regions. Together these techniques allowed the fate tracking of human CD34^+^ HSPCs to assess their lineage potential at a single-cell level.

The modified scATAC-seq protocol adds a fixation step and an adapted permeabilization step designed to retain mitochondria (MT) in cells for subsequent capture and library preparation (Figure 1A). This method achieved uniform coverage across the MT genome and greater than 30-50x coverage in a minimum of 1000 cells (Figure 1B-C) of the MT genome, which was suitable for variant calling and cellular “barcoding”. First, low-quality cells and MT alleles with low coverage and base-quality were filtered. Then additional variants were excluded using the Mitochondrial Genome Analysis Toolkit (MGATK), which removes variants with a low correlation of allelic reads across strands and a low variance-mean ratio (Figure S1A-B, see Methods).^28^ To de-multiplex each donor across conditions, the Vireo algorithm was applied on germline MT variants (Figures S1C-E, see Methods). The donor predicted variant allele-frequency (heteroplasty) revealed 29 variants of high mean VAF (mean >0.7 in donor, <0.1 in others), highlighting many donor-specific variants that were primarily transition mutations (Figure 1D). Performance was assessed by varying the number of donors and calculating Vireo’s likelihood score, the evidence lower bound. The “elbow rule” confirmed that the true number of donors was the inflection point in which performance gain was reduced when additional possible donors were added to the model (Figure S1D). These data demonstrate that mt-scATAC-Seq can be used to identify mitochondrial variants across the entire mitochondrial genome and to demultiplex mixed samples.

### Single CD34^+^ cells generate clonal populations of variable size identified by mitochondrial variants

Clone assignment is critical to track the fate of CD34^+^ stem cells at the single cell level. Somatic MT variants were used to facilitate clone assignment in cultures of human CD34^+^ cells (Figure 2A, Figure S1A), using a community-based k-nearest neighbors (KNN) clustering algorithm on heteroplasmy across single-cells (Figure S2).^28^ After removing clones with fewer than 5 cells, 26-50 clones per donor were detected per capture (Table S1). Clone size was variable between donors, with clone size in the input sample correlating with the clone size after 72h culture (Figure 2B,C). Clones were identified based on combinations of unique MT variants (Figure 2D, E). Clone assignment was based on heteroplasty (fraction of reads with the variant) rather than binary variant calls because the MT genome is heteroplasmic. Indeed, cells in clones were distinguished across a range of heteroplasmy (Figure 2A, 2E). Some variants were shared across clones, such as variant 5581G (Figure 2D,E), and these are predicted to have arisen from a common ancestral stem cell. Examination of the cumulative fraction of donor cells based on the percentage of total clones, ordered by size, indicated that the larger clones did not dominate the pool in these healthy donors (Figure 2F).

**Figure 1.**
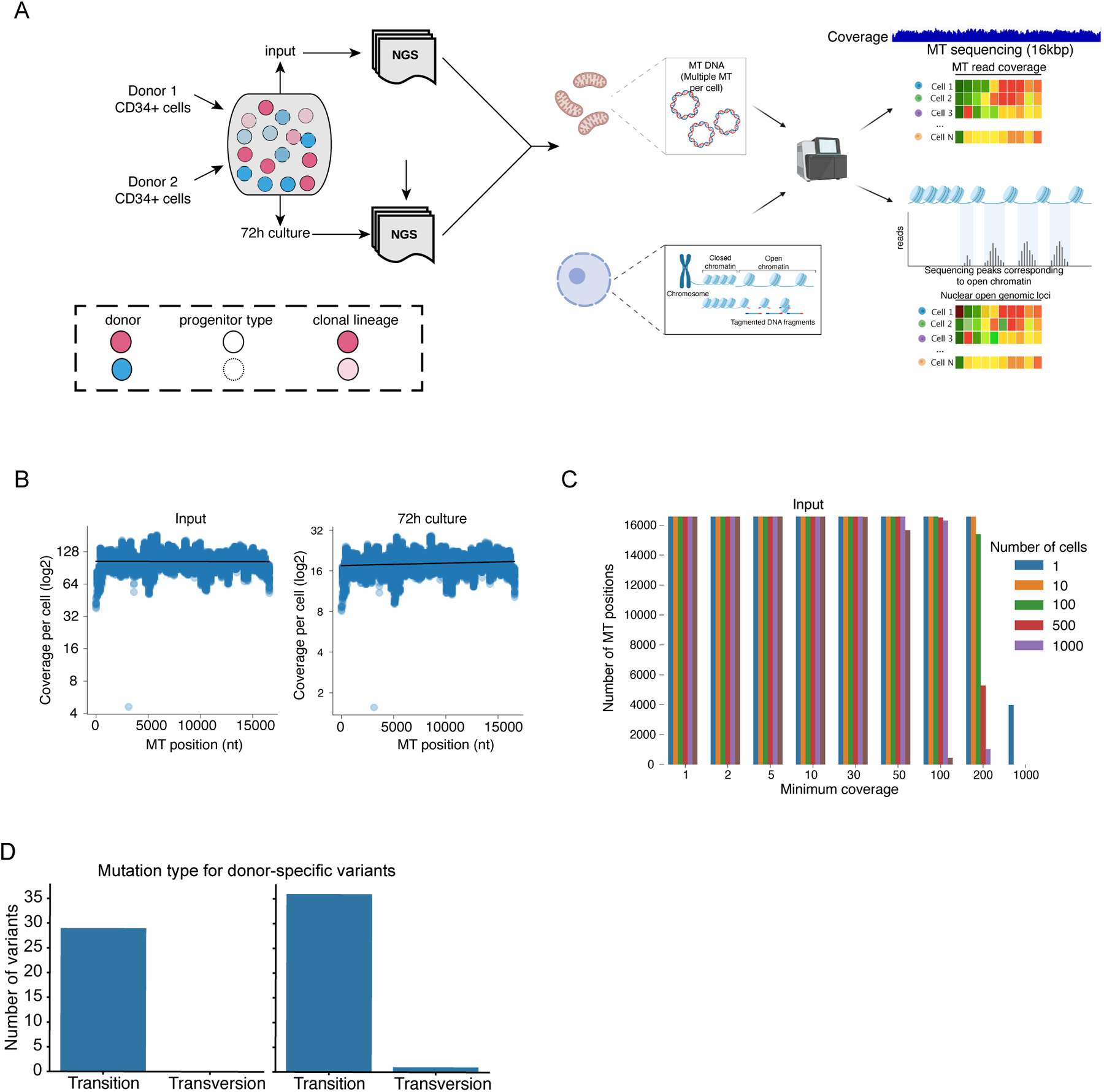
mt-scATAC-seq defines clonal lineages in mobilized human CD34+ cells. (A) Overview of mitochondrial mt-scATAC-seq workflow (B) Coverage across MT genome in single-cells across sequencing experiments. Black line is the mean at each position (C) Number of MT positions covered across a range of cells and coverage thresholds across batches. (D) Transitions and transversions detected in donor 1 (left) and 2 (right). Donor-specific variants defined as having greater than 0.8 VAF in over 90% of donor-assigned cells.

**Figure 2.**
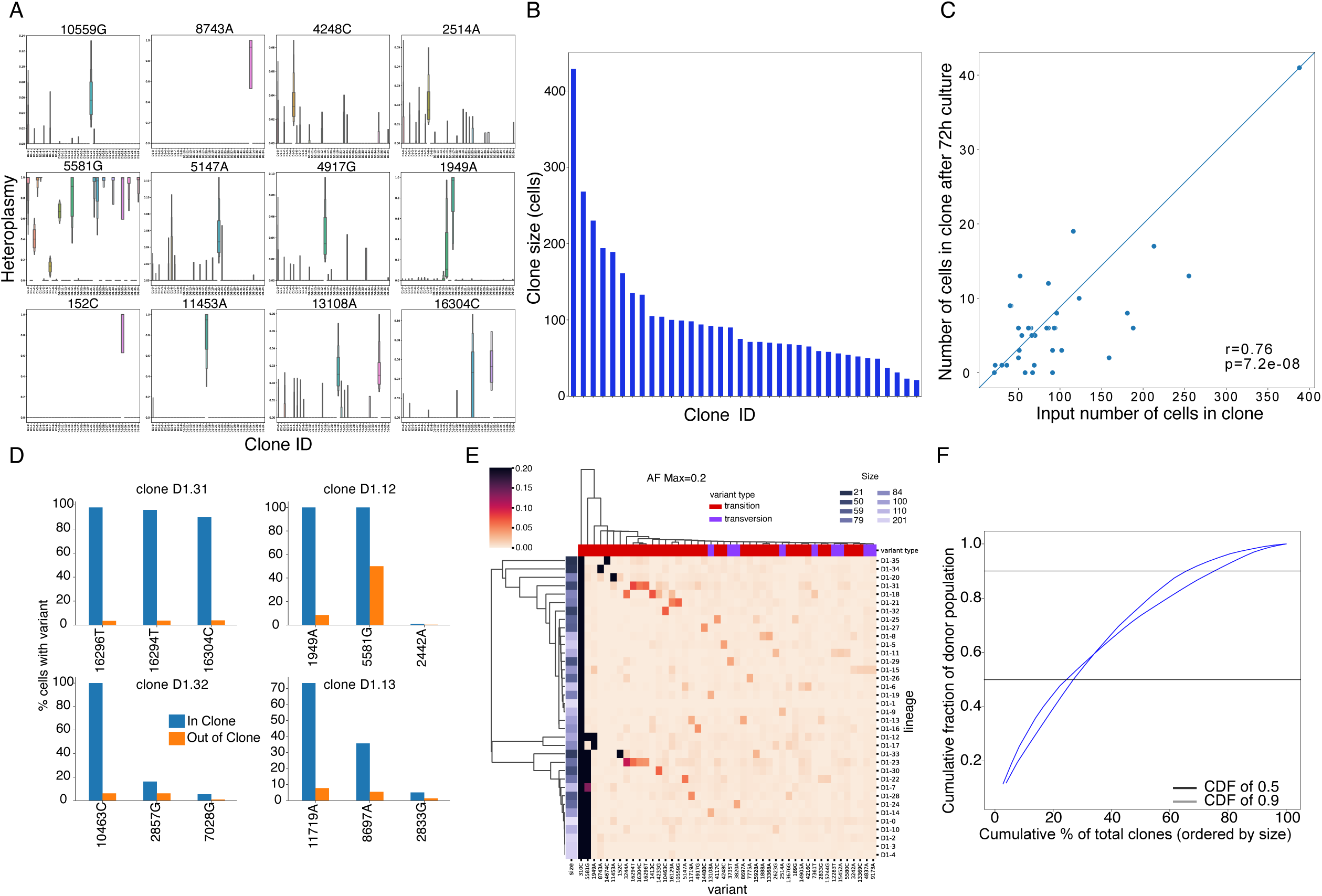
Single CD34+ stem cells expand into clones of variable sizes identified by mitochondrial variants. (A) Mitochondrial heteroplasmy in detected clones. Distribution of somatic mitochondrial barcodes (y-axis) across detected clones (x-axis) in Donor 1. Each variant is labelled as position followed by the alternative variant. Each boxplot is the heteroplasmy distribution of that allele in all cells within that clone. (B) Number of cells in each clone. (C) Scatterplot of number of cells across input and cultured cells. Pearson correlation r and p-value shown. (D) Barcodes detected in representative clones. In blue is the percent of cells with the barcode in the clone, and in orange is the percent in cells outside of the clone, shown for the top distinguishing barcodes for each clone. (E) Average heteroplasmy of each variant in each clone. Max cut-off at 0.2. Variant types shown for each variant and number of cells for each clone. (F) Cumulative distribution of the number of cells captured with increasing number of clones, sorted by largest to smallest, across both donors. Each donor line colored by the conditions used.

Different variant-calling and clone-detection workflows were then compared and we observed a high concordance of cell-pair clonal-relationships across parameters (see Methods). For example, we varied the number of neighbors (k) in the KNN algorithm and noticed a high concordance of clonal assignment with higher values of k. A lower performance was noted when the number of nearest neighbors in the KNN algorithm was low, as it resulted in larger, more sparse clusters, leading to lower consistency in clones detected when running the algorithm on a subsample of the population (Figure S2). While clonal detection using nucleotide barcoding is sensitive to the number of cells sampled and read depth, these smaller clones may represent a low fraction of total hematopoiesis.^25,29^ Taken together, these data demonstrate that mt-scATAC-Seq can be used to identify clonal lineages in human CD34^+^ cells using mitochondrial variants acting as natural barcodes. Additionally, the proliferative potential of CD34^+^ cells in *ex vivo* culture conditions is not related to the size of the clone within the CD34^+^ pool (Figure 2C, 2F).

### mt-scATAC-seq identifies lineage commitment of CD34^+^ cells in culture

After successfully mapping the clonal lineage of human CD34^+^ cells using the mitochondrial open-chromatin reads captured by mt-scATAC-seq, the nuclear genomic open-chromatin reads captured from the same experiments were then used to identify active genomic loci for determining cell lineage. Coupling MT and nuclear chromatin reads enabled the differentiation potential of each clonal population of CD34^+^ cells to be mapped. Nuclear open-chromatin reads were processed using conventional scATAC-Seq tools, enabling dimensionality reduction and cell clustering. Quality control of experiments showed a comparable number of detected peaks across experiments (Figure S5A-C). Data from input CD34^+^ cells and cultured CD34^+^ cells were embedded and clustered using uniform manifold approximation and projection (UMAP) and Louvain clustering. Twelve populations were identified using analysis of gene activity scores in exonic and promoter regions critical for each cell lineage (Figure 3A, Methods). As peaks were also found in intergenic and other non-coding regions (Figure S5D), transcription factor motifs were also identified in open-chromatin peaks using ChromVar.^30^ The gene activity scores of many lineage markers were differentially regulated between clusters supporting the differentiation of the human CD34^+^ cells to unique blood cell lineages, and also supporting the lineage-fate relationship of each cluster of these *ex vivo* cultured CD34^+^ human HSPCs (Figure S4). Transcription factor activity scores also supported the assignment of clusters towards the erythroid (GATA1-TAL1) and myeloid (SPI1 and CEBPE) lineages (Figure S4D). To improve the resolution of the lineage assignment by gene activity and transcription factor activity, cells were integrated from all samples (Figure S4). Subsequent analysis showed a higher resolution of clusters including two clusters in the neutrophil developmental lineage (Figure S4B). These data support the dual use of mt-scATAC-Seq for assigning clonal lineages with mitochondrial natural barcodes as well as nuclear accessible chromatin read signatures for assigning cell lineages.

**Figure 3.**
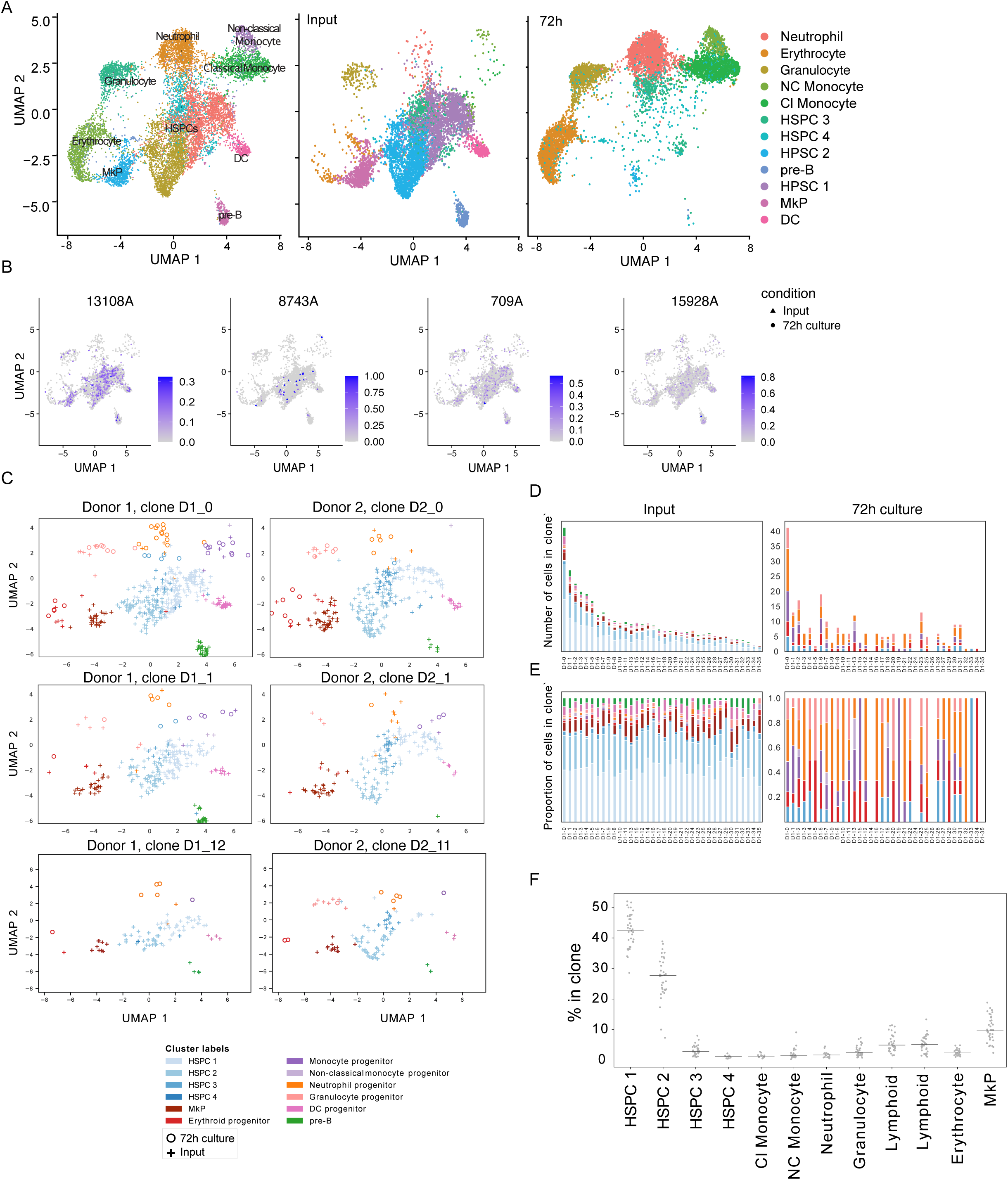
mt-scATAC-seq reveals minimum lineage-bias in the ex vivo differentiation of human CD34+ cells. (A) UMAP of cells in both input and culture colored by cluster. Left: Both conditions; middle: input; right: after 72 hour culture. (B) MT variants across UMAP for selected variants in donor 1. (C) Lineage fates of cells in input. Cells in representative clone in each donor embedded in UMAP. (D) Raw cell counts in each clone, colored by lineage cluster. (E) Percent of immune lineage clusters across each clone for donors. Color legend is same as in (A). (F) Percent of lineage in a clone, across all clones for donor 1 and donor 2.

### Lineage bias of human CD34^+^ cells using mt-scATAC-seq

The differentiation of CD34^+^ cells into diverse lineages in these assays indicated that an assessment of lineage bias could be performed on these clonal populations. When overlaying cells from clonal lineages and quantitatively analyzing their distribution, lineage-restricted cells were clustered with negligible evidence of lineage bias (Figure 3B-E & SF3E). To test if clones were biased to specific HSPC clusters, a hypergeometric test was applied, which detected few clone-cluster pairs with evidence of bias (Figure 3C-F). The clones D1-12, D2-11, D2-17, and D2-22 were overrepresented in lymphoid, granulocytes, and HSPC clusters (Benjamini-Hochberg adjusted p-value < 0.1, see Methods); however, these contained fewer cells overall. Furthermore, the lymphoid lineage cells were not found in the *ex vivo* differentiated condition (Figures 4 & S4). The lineage bias was also examined after pooling data from all donors using the normalized entropy metric (see Methods). Using this algorithm, if a clone is biased to one clone, it will display a lower entropy. Analysis of clones revealed little variability in clone entropy with smaller clones correlating with lower entropy (Figure S3F). These data suggest that stochastic changes in lineage commitment are involved, rather than biased differentiation of LT-HSC.

To further define the relationship between clone size, cell type, and lineage bias potential in a CD34^+^ clonal lineage, the lineage proportions in each clone were compared across cell types. This was also measured across all donors using an entropy measurement in each lineage and converting the clonal counts into probability distributions for each lineage. The lineage proportions in each clone varied widely across cell type (Figure 3 and S3). In contrast, the variation across clones within each lineage was smaller within each lineage compared to between the lineages (Figure 3F, S3F). While most clones showed no skewing across clusters, it was also necessary to determine if the resolution of clone detection influenced these data. To do this, individual MT variant barcodes across the HSPC clusters were examined and no significant biases across the lineage clusters in donor 1 and donor 2 were found (Figure 3B-F). Together, these data suggest that detectable HSPC clones have multi-potent capacity contributing to hematopoiesis in humans without substantial lineage bias.

To validate the results of mt-scATAC-seq, flow cytometry of CD34^+^ cell cultures was performed in parallel. Dimensionality reduction and automated clustering were performed on human CD34^+^ cells prior to culture and after 72h of culture using antibodies specific to human HSPC markers CD34/CD117 (c-Kit), lymphoid lineage markers CD3/CD19/CD56, granulocyte lineage markers CD66b/FcεRIα/Siglec8, monocyte lineage markers CD14/CD16/CD86/CD11c, and additional developmental and maturation markers CD45/CD10/CD101/CD11b/HLA-DR. Analysis of these flow cytometry data using dimensionality reduction identified 13 clusters including 4 HSPC subsets, 1 granulocyte-like cluster, 1 classical monocyte cluster, 1 non-classical monocyte cluster, 1 pre-B cluster, 1 DC cluster, 1 additional CD34^+^ cluster and 3 CD34^-^ clusters (Figure 4A, B). Consistent with mt-scATAC-seq, the pre-B and DC clusters were no longer represented in the cell population after 72h of culture, suggesting that these two populations may be contaminants of CD34^+^ cell sorting and undergo cell death in the *ex vivo* culture. The frequency of these clusters identified by mt-scATAC-seq and flow cytometry are consistent, supporting the validity of the mt-scATAC-seq approach and the capacity of the CD34^+^ cells to differentiate into diverse cell types in presence of SCF/IL-3/IL-6/Flt3L/G-CSF/GM-CSF (Figure 4C). These flow cytometry data support the *ex vivo* mt-scATAC-Seq mitochondrial barcoding system to study short-term differentiation of human CD34^+^ cells to lineage-committed cells.

**Figure 4.**
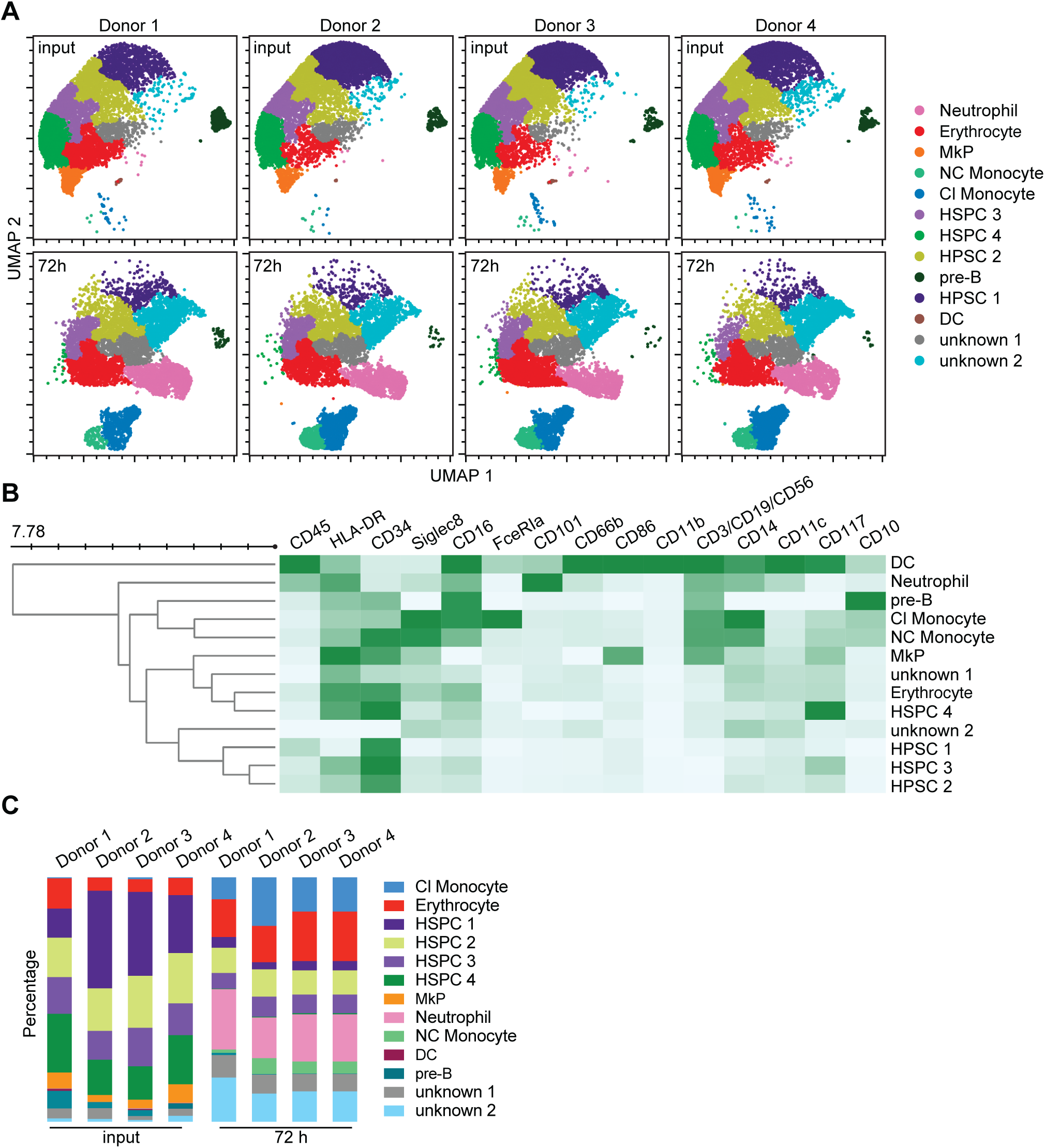
Flow cytometry identifies variable cell lineages in human CD34+ stem cell steady-state and in ex vivo culture. (A) UMAP shows cell clusters identified by CD45, HLA-DR, CD34, Siglec8, CD16, FceRIa, CD101, CD66b, CD86, CD11b, CD3, CD19, CD56, CD14, CD11c, CD117, CD10. (B) Heatmap shows expression levels of indicated markers in each cell cluster. (C) Bar graphs show the proportion of each cluster in each donor before- and after-72h culture.

### Assessing lineage bias in differentiation of single mouse HSC in vivo

To compare the differentiation of clonal human CD34^+^ HSC, as assessed by mt-scATAC-seq, and mouse LT-HSC, a four-step method for mouse LT-HSC purification and transplantation was optimized and used to evaluate long-term engraftment and differentiation of single LT-HSC (Figure 5A). To confirm that the phenotype of single cells was consistent with HSC, reconstitution in primary and secondary lethally-irradiated recipients was examined by multicolor flow cytometry panels that allowed for the identification of donor white blood cells (CD45.2) from recipient (CD45.1/CD45.2) and support (CD45.1) cells (Figure S7). The conditions used to isolate HSC yielded >1% engraftment of 12/18 recipient mice transplanted with a single HSC, and 100% of recipient mice transplanted with 10 HSC or 100 HSC (Figure 5B-D). Changes in developmental-related reconstitution and ratios of myeloid and lymphoid cells was evident between 1 month and 4 months post reconstitution (Figure 5B). Flow cytometric analysis of donor myeloid and lymphoid populations in bone marrow, spleen, and blood, and across time in the peripheral blood, indicated that single LT-HSC were capable of high efficiency long-term multi-lineage repopulation. These data demonstrate that this sorting strategy results in high efficiency engraftment and reconstitution of the hematopoietic system.

When compared to wild-type steady state (not transplanted) mice, flow cytometric analysis of hematopoietic tissues from irradiated single cell HSC recipient animals revealed substantial variability in reconstitution of cells from the myeloid and lymphoid lineage, pointing to potential lineage bias of LT-HSC used for reconstitution (Figure 5D). However, this variability in lineage reconstitution was also evident when 10 or 100 HSC were used to reconstitute animals. In these latter cases, any bias would be predicted to be masked by the average response of the larger numbers of LT-HSC population in these animals. Taken together, the variable reconstitution in the myeloid and lymphoid lineage suggests that other factors, such as the inflammatory milieu and the hematopoietic niche of the lethally-irradiated host, must also contribute substantially to the behavior of donor LT-HSC.

## Discussion

Lineage-bias in the hematopoietic stem cell pool has been widely investigated *in vivo* but relies on a number of key assumptions: (1) the donor cell population alleged to be HSC are not multi-potent progenitors or lineage-committed progenitors; (2) *in vivo* barcoding and other labeling methodologies are labeling hematopoietic stem cells with high efficiency; (3) that variation in the repopulation of myeloid and lymphoid cells in recipient animals reconstituted with single HSC is not influenced by irradiated host cells or support bone marrow donor cells such as Th1 cells,

Foxp3+ regulatory T cells and NK cells; (4) that skewed differentiation is not a consequence of pressures exerted on multipotent progenitors and/or lineage-committed progenitors; (5) that transplantation of typically >50 HSC can be relied on to investigate the cell-intrinsic lineage-bias of individual hematopoietic stem cells; (6) that the dynamic changes in ratio of myeloid:lymphoid:erythroid following transplantation are acknowledged; and (7) that dwindling proportions of donor hematopoietic cells over time is consistent with the behavior/function of HSC at steady state.^31–35^

**Figure 5.**
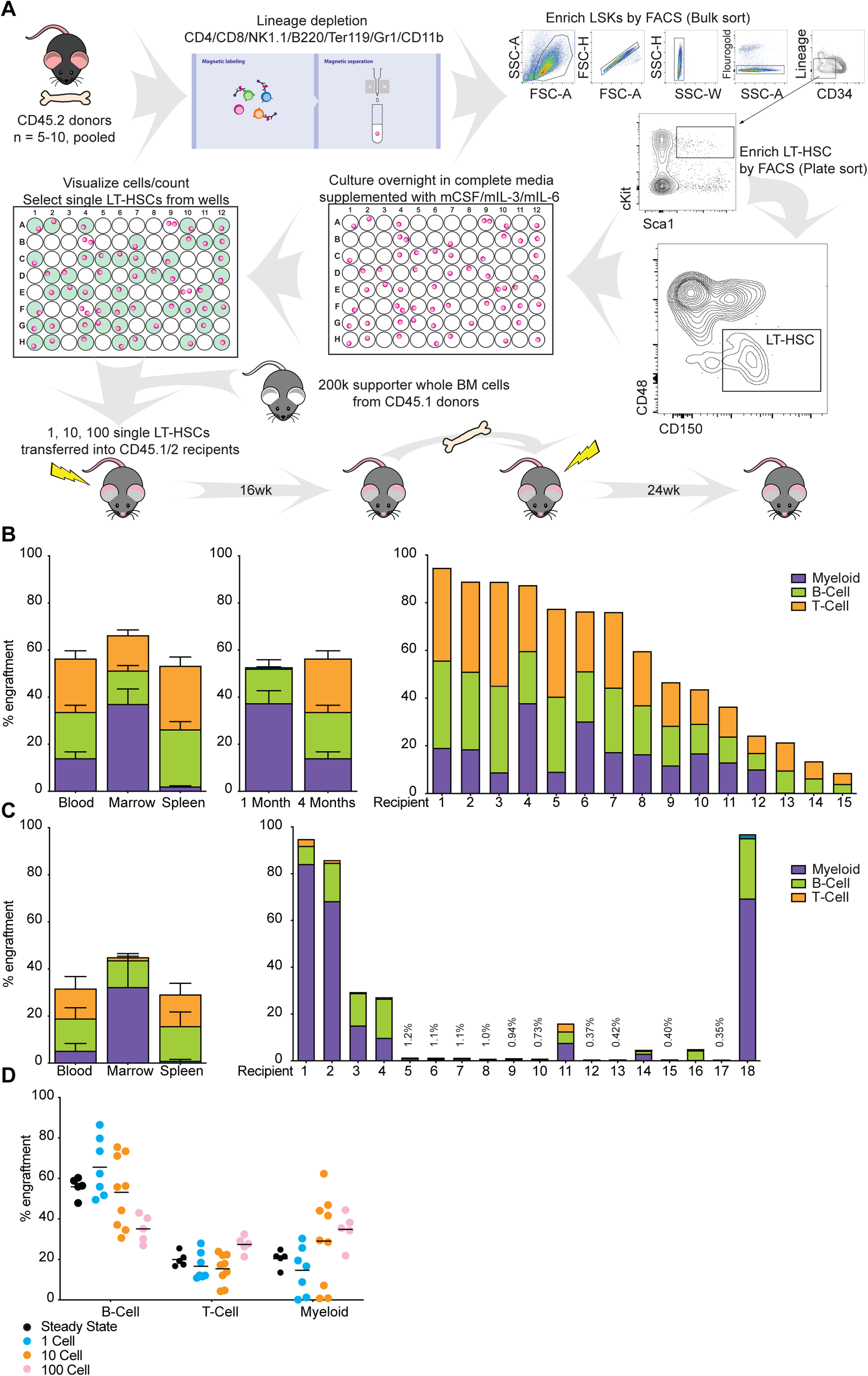
LT-HSC reconstitution varies by recipient, tissue, and time post-engraftment. (A) Four-step method for purification and transplantation of LT-HSC to evaluate long-term engraftment and differentiation. Step 1: Bone marrow from 5-10 CD45.2 wild type mice was pooled and lineage depletion was performed using magnetic columns with a custom lineage antibody cocktail. Output cells were stained for flow cytometry sorting. Step 2: CD34-Lineage¬-Sca+Kit+ (CD34-LSK) cells were bulk sorted. Step 3: LT-HSC (LSK CD150+CD48-) were plate sorted into 96 well plates. Step 4: Cells were cultured for 12-18 hours in complete media supplemented with mCSF/mIL-3/mIL-6. LT-HSC were mixed with whole bone marrow from CD45.1 wild type mice and injected into lethally-irradiated recipient mice. (B) Reconstitution of peripheral blood, bone marrow, and spleen at 1 and 4 months post-transplant of lethally-irradiated mice with 10 LT-HSC. Left panel: 4 months post-transplant. Middle panel: peripheral blood 1 month and 4 months post-transplant. Right panel: Reconstitution of peripheral blood from individual mice. (C) Reconstitution of peripheral blood 4 months post-transplant of lethally-irradiated mice with 1 LT-HSC. 12/18 mice engraft with greater than 1% donor LT-HSC-derived cells in myeloid and lymphoid compartments. (D) Comparison of lineage reconstitution in the blood of successfully-engrafted lethally-irradiated mice receiving 1, 10, or 100 LT-HSC compared to non-irradiated wild-type mice at steady state.

In this study, significant variation in the ratio of myeloid:lymphoid cells in mice reconstituted with a single HSC is demonstrated. This could be interpreted as evidence of lineage-bias in donor HSC; however, the myeloid:lymphoid ratio in mice reconstituted with 10 HSC shows similar variation in the myeloid:lymphoid ratio. Critically, mice fully reconstituted using 100 HSC is not equivalent to steady state, or mice reconstituted with 1 HSC or 10 HSC. If lineage bias exists between HSC, then the variation in overall myeloid:lymphoid ratio should reduce with increasing numbers of donor HSC as the average output of lineage-biased HSC approaches that of mice reconstituted with whole bone marrow or compared to mice at steady state. However, a reduction in the variation of myeloid:lymphoid ratio was not observed in mice receiving 10 HSC despite an increase in successful engraftment and efficiency of reconstitution in these experiments. These findings argue against substantial lineage bias in hematopoietic stem cells capable of reconstituting mouse hematopoiesis. Rather, it suggests that environmental and inflammatory stimuli may perturb differentiation of multipotent progenitors and lineage-restricted hematopoietic progenitor cells in the irradiated host. These findings are consistent with recent fate mapping studies from fetal liver cells which conclude that the expansion potential of HSC is not pre-determined but rather influenced by the niche.^26^ Similarly, human gene therapy studies also concluded that HSC distributed their clonal progeny without lineage bias.^27^

The methodology also provides an opportunity to enhance the study of mouse and human HSC *ex vivo* and *in vivo*. With improved, standardized, and concurrently validated, methods for HSC isolation and HSC lineage tracing, the study of genetic and biochemical regulators of HSC proliferation and differentiation can be explored using diverse methodology including clonal culture assays, single cell transcriptomics and epigenomics, and *in vivo* differentiation and function^33^. The effects of cytokines and microbial products on gene expression, cell function, and donor cell engraftment can also be explored with higher resolution and confidence. Barcoding and transplantation techniques provide excellent clone-detection methods; however, the cellular and immune response to these editing techniques can introduce confounding effects that require consideration in experimental and therapeutic design^36^, a factor that is avoided using naturally occurring mitochondrial DNA barcodes.^28,37–39^ The rapid advances in novel conditioning regimens and genetically-modified recipient mice with an enhanced capacity for engraftment demands improvements in reproducible methods for the isolation of pure HSC populations.^34,35^ Comparison of the engraftment efficiency of single HSC in lethally-irradiated mice and in non-hemoablative murine models is also needed. Perhaps most relevant for the field of hematopoiesis would be a unified approach with unambiguous functional definitions and not simply molecular definitions when reporting the outcome of experiments that investigate hematopoietic stem cells and hematopoietic progenitor cells.

## Acknowledgements

This work was supported by NIH Grant R01AI155869, HL152958, R35GM119850, the V Foundation for Cancer Research, and a T32 (5T32HL007574-36 to MBM).

## Methods

### Human primary CD34+ cells

Cryopreserved CD34^+^ hematopoietic stem and progenitor cells were obtained from Gilead (Donors 1-4, Donors A-D for FACS sorting) or StemCell Technologies (Donors 5-8). Adult donors gave informed consent for the collection of CD34+ cells. Samples, where applicable, were cultured for 72h in DMEM/10%FBS/10%CO_2_ and a cytokine culture consisting of 100 ng/mL recombinant human SCF/IL-3/IL-6/Flt3L/G-CSF/GM-CSF. The CD34^+^ samples were de-identified and processed in both the mt-scATAC-seq library preparation and FACS sorting.

### mtscATAC-seq library preparation

mt-scATAC-seq libraries were generated by adapting the 10X Genomics protocol for single cell ATAC seq, according to modifications made by Lareau et al^28^. This modified protocol exploits a fixation step and a modified permeabilization protocol to retain mitochondria in cells for subsequent capture and library preparation (Figure 1A). Whole cells were retained following an adapted protocol of 10X Genomics Single-cell ATAC-seq. Cells were fixed in 1% formaldehyde and both digitonin and Tween 20 omitted in the lysis and wash buffers to generate a higher retention of mitochondrial DNA fragments per single cell. Cells were washed twice in PBS and centrifuged for 5 min at 400xg in 4°C. Samples were multiplexed by taking the same number of cells per patient sample to get a total number of cells above 10^5^ to account for cell loss. Samples were incubated in lysis buffer for exactly 3 min on ice prior to washing. After centrifugation of sample at 500xg for 5 min, the cell supernatant was discarded, and the cell pellet was resuspended in 1X Diluted Nuclei buffer. Cells were processed according to the Chromium Single Cell ATAC Solution user guide using the Chromium Next GEM Single Cell ATAC Library Kit, Chromium Next GEM Single Cell ATAC Gel Bead Kit, Chromium Next GEM Chip H Single Cell Kit, and Single Index Kit N Set A. Lastly, quality control (QC) tests were run on each library prep using Agilent TapeStation High Sensitivity D1000 (Agilent) and Qubit dsDNA HS Assay kit (Invitrogen) prior to sequencing.

### Processing of mt-scATAC-seq sequencing fragments

Processing of mt-scATAC-seq reads was performed as previously reported^28^. The cellranger-atac count command from cellranger v6.1.1 was used to generate bam, peak genomic regions and peak-fragment count files. The hg38 reference genome was modified by hard-masking nuclear regions that align to the MT genome with single bp errors^28^ (regions taken from https://github.com/caleblareau/mitoblacklist/tree/master/combinedBlacklist). Reads were trimmed to remove the adapter and primer sequences, and then aligned using BWA-MEM^40^. Open-chromatin peaks were detected, and cell barcodes were filtered. For peak-calling, reads were aggregated across all cells to boost signal, and a global threshold was applied to select candidate regions above background genomic noise. This was done by fitting negative-binomial distributions to estimate background and peak likelihood in the candidate regions. Local-maxima peaks within this region were then found and a local threshold was applied, generating peaks of various sizes. For cell-calling, potential barcode multiplets were collapsed by masking the minor barcode, and barcodes were removed using a threshold for fraction of fragments in the peak using a mixture model of two negative binomial distributions to capture the signal and noise, with an odds ratio threshold of 100000.

### Variant calling in the MT genome

Cells were filtered with less than 200 bp in the MT genome and fragment duplicates removed. Positions were removed with less than ten cells with at least 50x coverage, and with less than 10 cells having 5x coverage of a putative variant at that position. Additionally, cells required an average Phred base quality score (BQ) of over 20 at the putative variant. MGATK filters removed variants with low strand concordance and low variance-mean ratio for each variant across all cells in a sample. The thresholds used were the same as previously reported^28^, with concordance of 0.65 and log 10 variance-mean ratio of -2.

### Separating multiplexed donor cells

To separate donors from the same sequencing run, the algorithm Vireo^41^ was used, which is a variational Bayesian inference algorithm that reconstructs each donor’s allele frequency profile (the donor’s mean allele frequency is the latent variable) and assigns a probability of each cell to that donor. Any cell with less than 0.9 probability to be assigned to a clone was removed. The algorithm also assigns a ‘doublet’ probability for each cell, which is the likelihood of the cell being part of multiple donors versus one. Cells with more than 0.1 probability of a doublet were also removed. To ensure the donors called were correct, the number of donors in Vireo +/-2 from the true number of donors was examined. The model’s reconstruction likelihood score, the evidence lower bound (ELBO), used in variational autoencoders, is saved for each donor parameter, and the ‘elbow rule’ is then used, which finds the error’s inflection point upon increasing the number of donors. Donor specific homozygous variants were calculated as having a mean allele frequency greater than 0.9. In all our cases, the true number of donors is where the elbow occurs.

### Clonal detection using MT barcodes

After computationally separating the donors, the single-cell variant allele frequency was calculated for high-coverage positions to reduce spurious clone-calling, and then MGATK was performed providing a new set of called variants for each donor. To detect cells of the same clone, the k-nearest-neighbors Leiden-based community detection algorithm was used^42^. The resolution parameter was set to 30, after assessing values of 30-50, and the cosine distance cutoff of the algorithm was set to 3.5. To measure consistency across workflows, cell pairs were examined to determine if they were either assigned to the same clone in both methods, assigned different clones in both, or assigned the same clone in one method but not the other (negative samples). We compared the fraction of the cell pairs that overlapped with each other (Figure S28). In Figure S2B, cell population was subsampled, and an adjusted normalized mutual information score was calculated between the cell-clone assignment in the sampled clone composition and the full sample detected clones.

To calculate the percentage of cells with the barcode in a clone and outside a clone in Figure 2D, variants were binarized with a minimum of 2 reads and an allele frequency of 0.001. The top 3 variants with the highest positive difference in percentage between clones and non-clones was chosen. For Figure 2E, complete-linkage using cosine similarity was used, setting allele frequency of >0.2 to 0.2 to improve visibility. Barcodes with an average of less than 0.01 in each clone were removed. In Figure 2A, the distribution of each barcode was plotted across cells in each clone using a boxenplot with default parameters in seaborn v0.11.2, which is a modified form of a boxplot that better represents the distribution for large data (https://github.com/heike/stat590f).

### Processing single-cell nuclear open-chromatin regions

To examine the peaks detected using the nuclear open-chromatin reads in each cell, the Signac (V1.4) protocol was used to integrate conditions, preprocess, and binarize the cells, run latent-semantic indexing (LSI), followed by UMAP dimensionality reduction, and KNN Louvain clustering to assign cluster labels^43^. Integration was done by comparing input and cultured cells or by integrating all sequencing runs.

To examine open-chromatin regions and aggregate data across experimental runs, the detected peaks were merged by expanding the peaks with overlap across runs. Peaks < 20 bp and >10,000 bp were removed and fragment counts were re-computed. A Signac model was used to remove regions with < 10 cells, and cells with < 200 features. Additionally, data were filtered by keeping peaks with: a) ≥ 10 and < 15,000 fragments; and b) ≥15% of the nucleotides in reads found in the peak was also covered in the peak (since a read can span the peak region and outside the region). Cells were also retained: a) with a nucleosome signal of ≥4 (i.e. the ratio of mononucleosomal to nucleosome-free fragments per cell); b) with a TSS enrichment of ≥ 0.2 (as defined previously); and c) with a ratio of reads aligned to blacklist regions over reads aligned to peaks < 0.05.

Peaks were binarized and a term frequency–inverse document frequency (TF-IDF) was assessed followed by SVD, which combined is the latent-semantic indexing method. UMAP was then run on dimensions 2-50, as the first factor correlates with depth. After this, runs were integrated using *FindIntegrationAnchors* of the Seurat package using the lsi transformed data^43^. After integration, UMAP on dimensions 2 to 30 of the integrated lsi components was utilized, then clustered using *FindNeighbors* and *FindClusters* with the SLM algorithm^44^.

### Annotating cell clusters using lineage markers

Cells were annotated by taking known lineage markers of both gene activity and TF activity and overlaying the density of the feature across the UMAP embedding. Gene activity scores for each gene was calculated by summing the number of peaks found in a gene and 2 kb upstream. Feature counts for each cell are divided by the total counts for that cell, multiplied by the median gene activity in that cell, and then natural log transformed to obtain an activity score. TF activity was calculated using the chromVAR extension in Signac, which estimates activity based on the number of TF motifs detected in a cell’s open-chromatin peaks^28^. Manual annotation was performed on the clusters using both the gene and TF activity in known markers.

### Hypergeometric test to measure lineage bias in clones

To detect clonal bias towards a specific lineage, a hypergeometric cumulative distribution test was used for each clone-cluster pair, and p-values were adjusted using the Benjamini-Hochberg method to control the false discovery rate. A significance threshold of 0.1 was used, but to account for clone and cluster sizes affecting the test, a non-parametric null distribution was created in which the cluster labels for each cell were shuffled 1000 times and the p-values for each clone-cluster pair computed. The p-values in each simulation were used as a background distribution, and empirical p-values were calculated for each clone-cluster pair, a significance of p=0.1 was used in reporting significance values.

### Clone and lineage entropy measures

To measure the lineage-bias across clones in Figure S3F, a normalized entropy metric was used. The ‘HSPC’ lineage clusters were removed, and the frequency of each cell type was assessed in each clone, and then used as the probability distribution. The standard entropy measure was calculated using entropy from the SciPy v1.7.3 stats package^45^, and was normalized to a value between 0 and 1 by dividing by the natural log of the number of clones.

### Flow-cytometry for human CD34^+^ cell cultures

Human: flow-cytometry was done for four healthy CD34+ donors, and culturing was done as mentioned above. Staining was performed in FACS buffer (D-PBS + 1% human serum + 0.1% sodium azide + 2mM EDTA) on ice. Cells were filtered through sterile 70 μm cell strainers to obtain a single cell suspension. Prior to staining, human Fc receptors blocking reagent (Biolegend) was added for 15 min. Staining was performed for 30 minutes in a final volume of 100ul. Cells acquired using a LSR Fortessa (BD Biosciences). All flow cytometry analysis performed on live cells. The markers used for dimensionality reduction in were HLA-DR, CD117, CD11c, CD11b, CD34, CD10, CD45, CD86, FcεRIα, CD16, CD14, CD66b, CD101, Siglec8, CD3, CD19, CD56.

### Four-step method for murine LT-HSC purification and transplantation

To evaluate long-term engraftment and differentiation of LT-HSC a four-step method for purification and transplantation of LT-HSC was used. First, bone marrow was pooled from the femur and tibial bones of 5-10 male and female 6-12 week old C57BL/6J mice (expressing CD45.2) and red blood cells lysed. Lineage depletion was performed using magnetic columns (Miltenyi) with a custom biotinylated lineage antibody cocktail (CD8a, CD4, CD11b, Gr1, Ter119, B220; BioLegend) with streptavidin magnetic beads (Miltenyi) [Step One]. Output cells were stained for FACS sorting with Lineage, CD4, CD8a, CD34, CD48, CD150, CD117/cKit, Sca1 (BioLegend), and Fluorogold (Sigma). Bone marrow was bulk sorted [Step Two] for Lineage^-^Sca-1^+^cKit+ (CD34^-^ LSK) on a BD FACS Aria II. The output of the first sort was then plate sorted for 2, 10, or 100 LT-HSC (LSK CD150^+^CD48^-^) into 96 well plates [Step Three]. Two LT-HSC were sorted instead of targeting one LT-HSC to account for dead volume in the syringe during injection and thereby eliminating avoidable failures of engraftment. Cells were then cultured for 12-18h in DMEM/10%FBS supplemented with 100ng/mL mSCF/mIL-3/mIL-6 (BioLegend) at 37°C at 5% CO_2_ [Step Four]. The following day, cells were visualized on a microscope before transplantation. LT-HSC were mixed with 2x10^5^ whole bone marrow from C57BL/6J CD45.1 wild type mice and injected retro-orbitally into lethally irradiated (2 x 550R) recipient mice (CD45.1/CD45.2).

## Code availability

All code used for data processing and analysis for this study has been deposited here, where it will be made publicly available upon acceptance of this work: https://github.com/LewisLabUCSD/Mito_Trace

---

**Figure S1.**
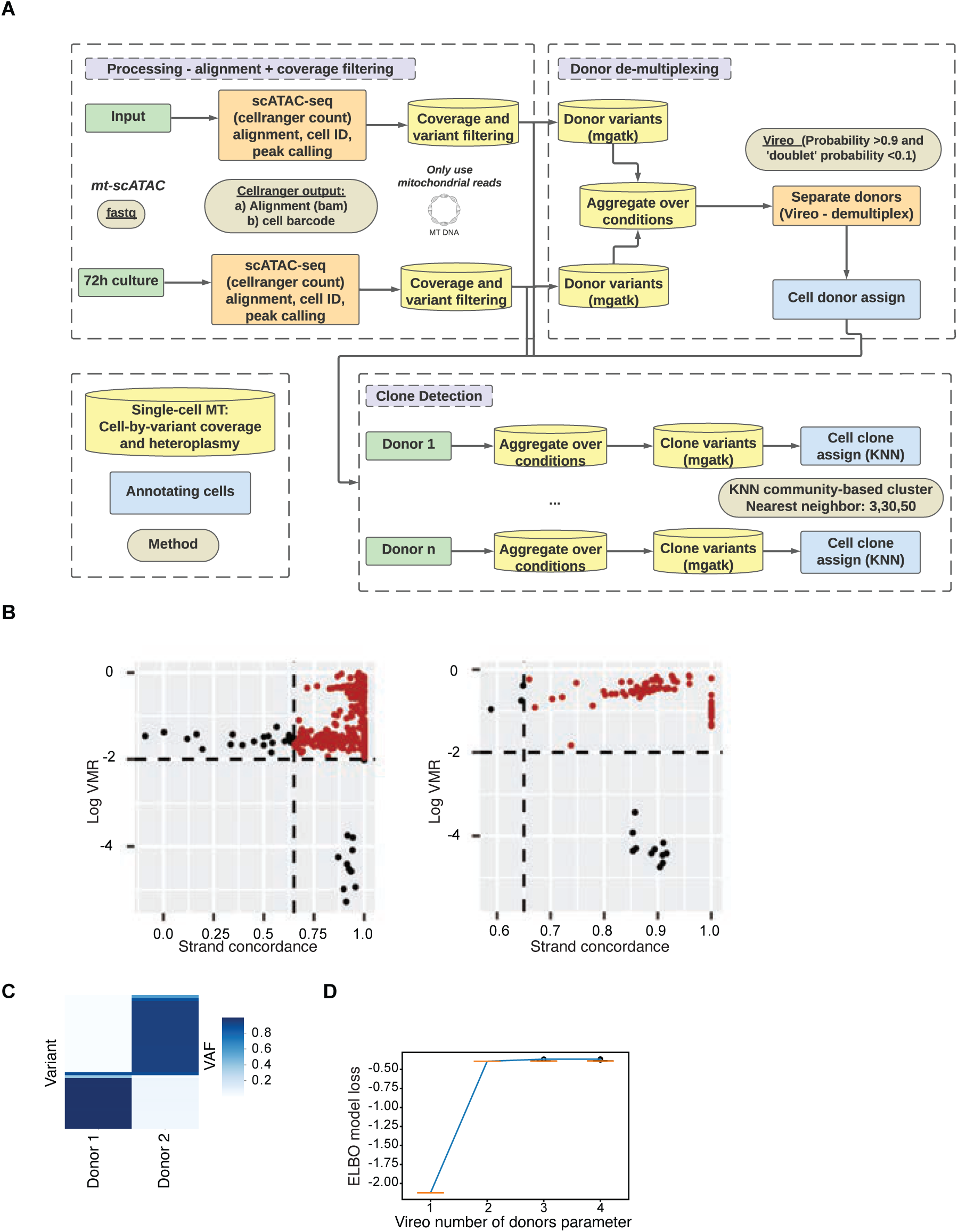
Pipeline and de-multiplexing in mt-scATAC-Seq experiments. (A) NGS processing, donor de-multiplexing and clone detection workflow. (B) MGATK algorithm used to call variants in the MT genome. Each point is a variant, and variants colored red pass the variance-mean ratio (VMR) and strand concordance thresholds. Left panel: input cells; right panel: 72 h culture. (C) Donor mean allele frequency. (D) The number of clusters (i.e. the number of donors) was varied and the Vireo likelihood score, the evidence lower bound (ELBO) was calculated. The “elbow rule” was then used to confirm that the true number of donors (n=2) was the inflection point in which performance gain was reduced when additional possible donors were added to the model.

**Figure S2.**
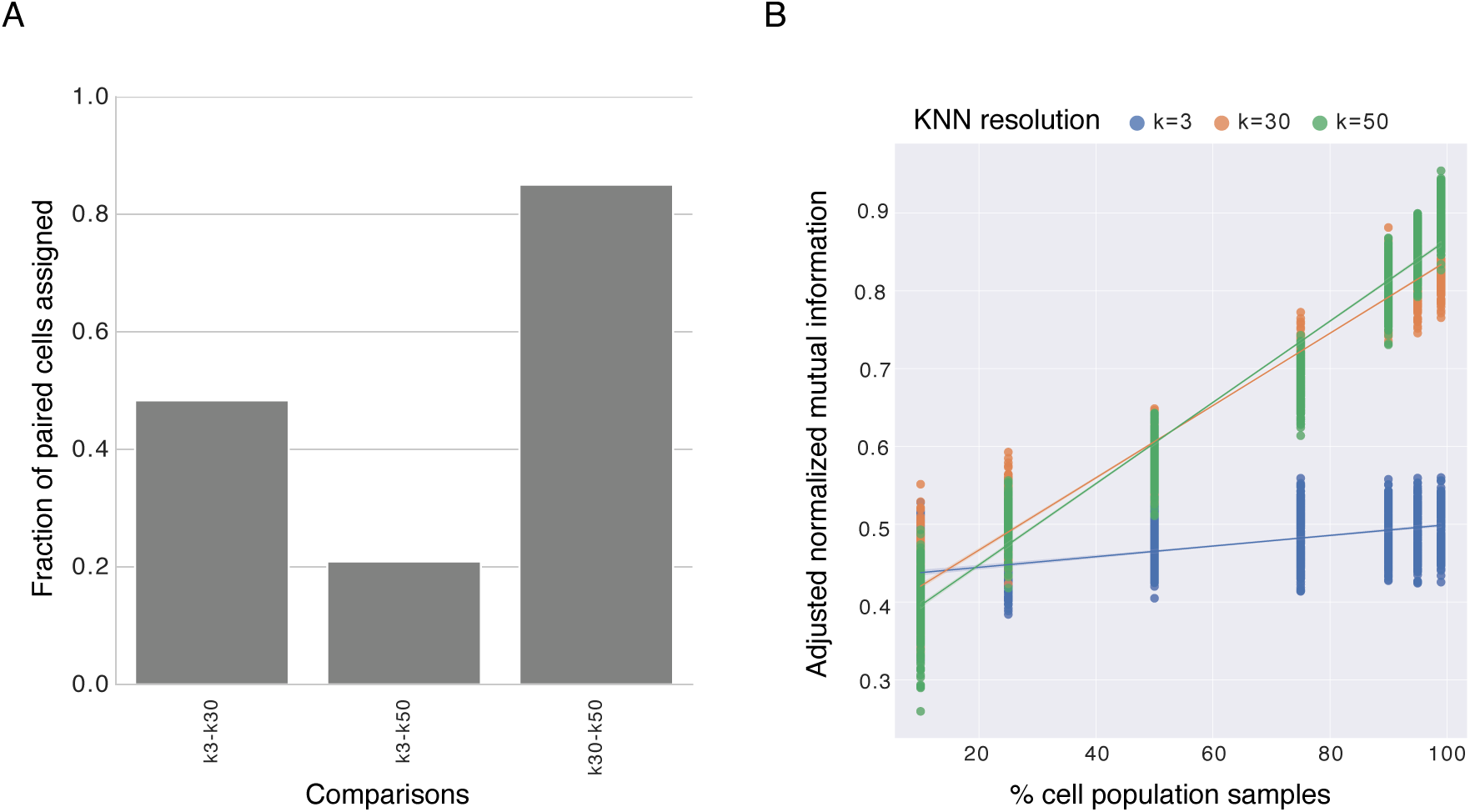
Clone assignment in mt-scATAC-Seq experiments. (A) Clone assignment comparison. K nearest neighbor of 3, 30, and 50 were compared by finding the number of cell pairs that are either in the same clone or in different clones in both methods. The score is then normalized to the fraction of paired cells assigned to the same clone across KNN resolution. (B) Subsampling cells from 10-99% 1000 times, calculating adjusted normalized mutual information between clones in sub-sampled run and the full population.

**Figure S3.**
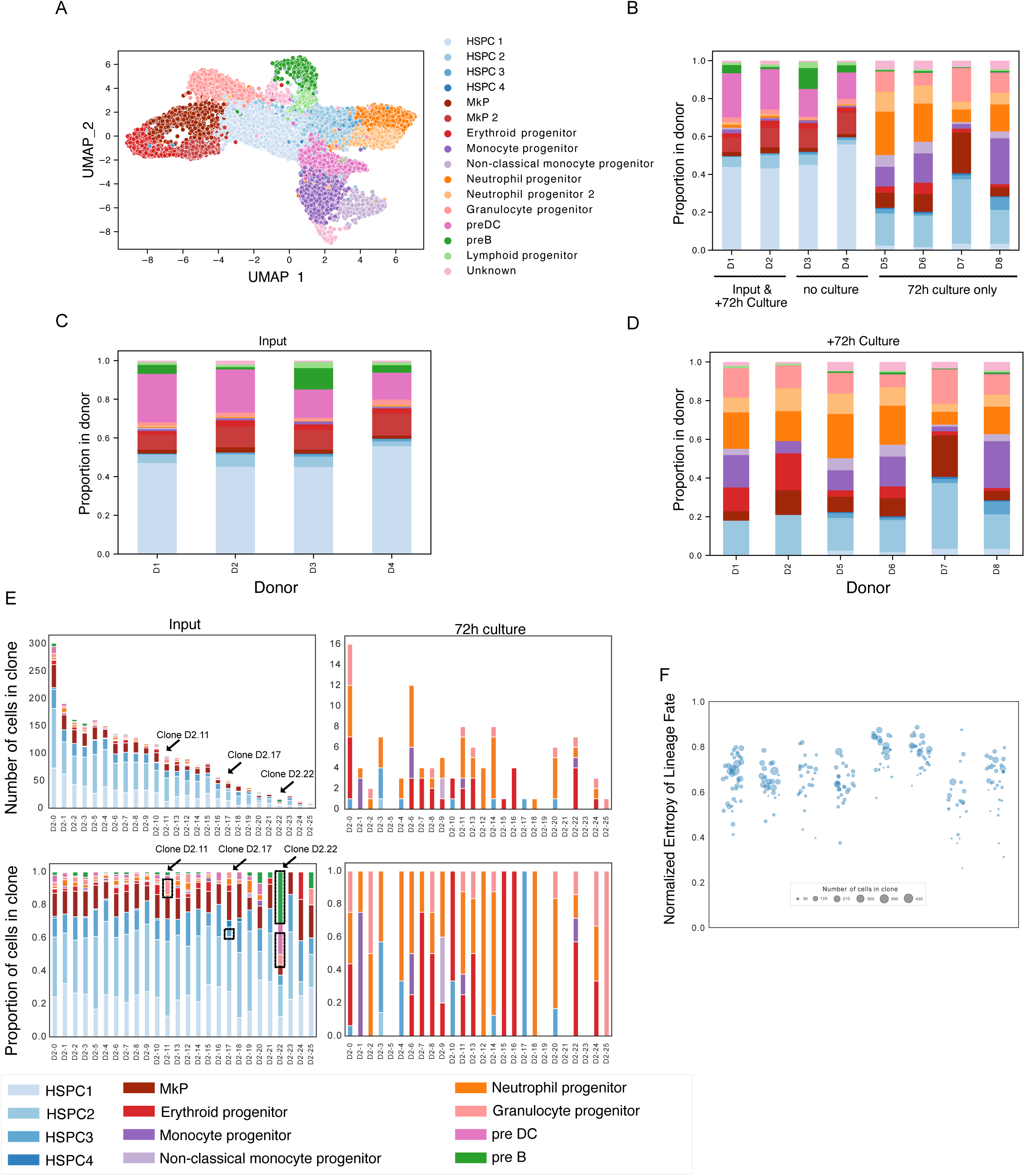
Minimum lineage-bias in human CD34+ HSPC clones across all donors. (A) Distribution of cells across all donors (n=8) and conditions on UMAP, colored by annotated cluster labels (B-D) Proportion of cells across HSPC clusters in each cell population studied, both input CD34+ cells and in cells cultured for 72h. (E) Normalized entropy of lineage fate in each clone after 72h culture, sorted by rank within each donor.

**Figure S4.**
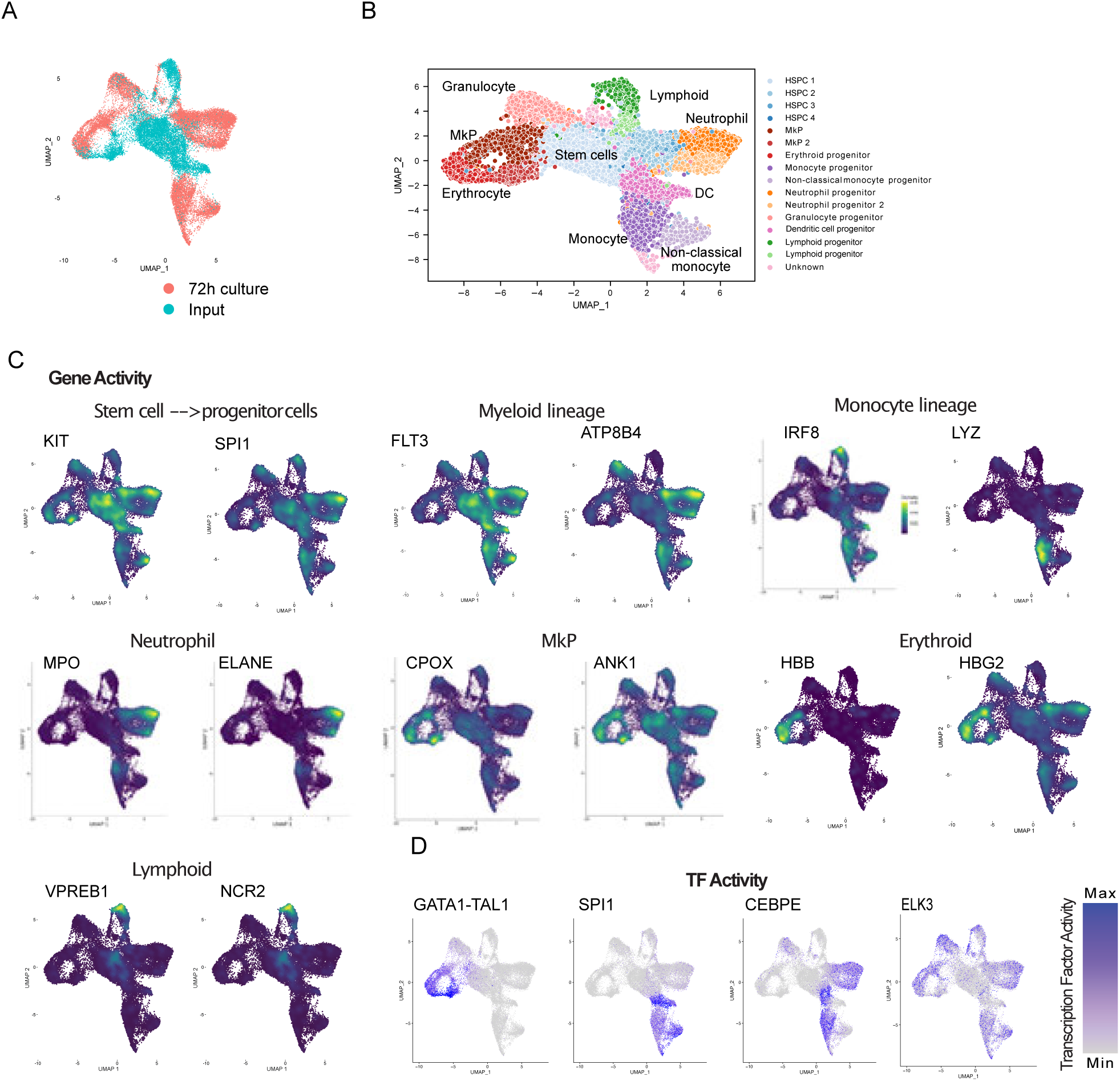
Characterizing cells by lineage markers in nuclear open-chromatin peaks across all donors. (A) Each batch sequencing run overlaid on UMAP. (B) Cells colored by clusters detected by Seurat’s SNN method, and UMAP annotation labels overlaid on UMAP. (C) Gene activity scores for select markers overlaid on UMAP. (D) Transcription factor activity scores for select markers.

**Figure S5.**
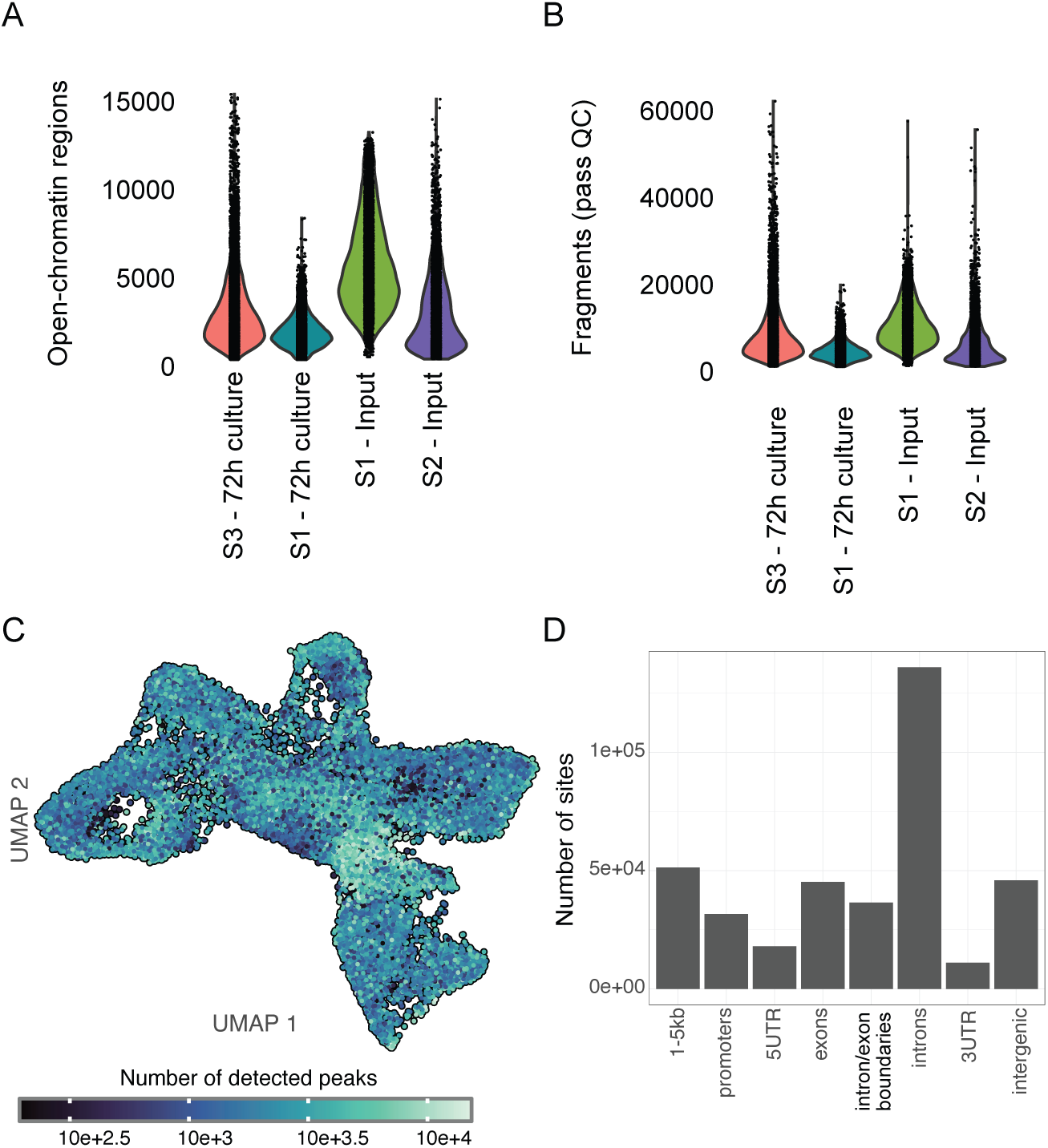
Detecting nuclear open-chromatin peaks. (A) Number of detected open-chromatin peak regions per cell. (B) Number of fragments that are non-duplicated and pass QC filters (see Methods). (C) UMAP illustrating the number of peaks detected for each cell. (D) The genomic location of each peak for each open-chromatin site detected.

**Figure S6.**
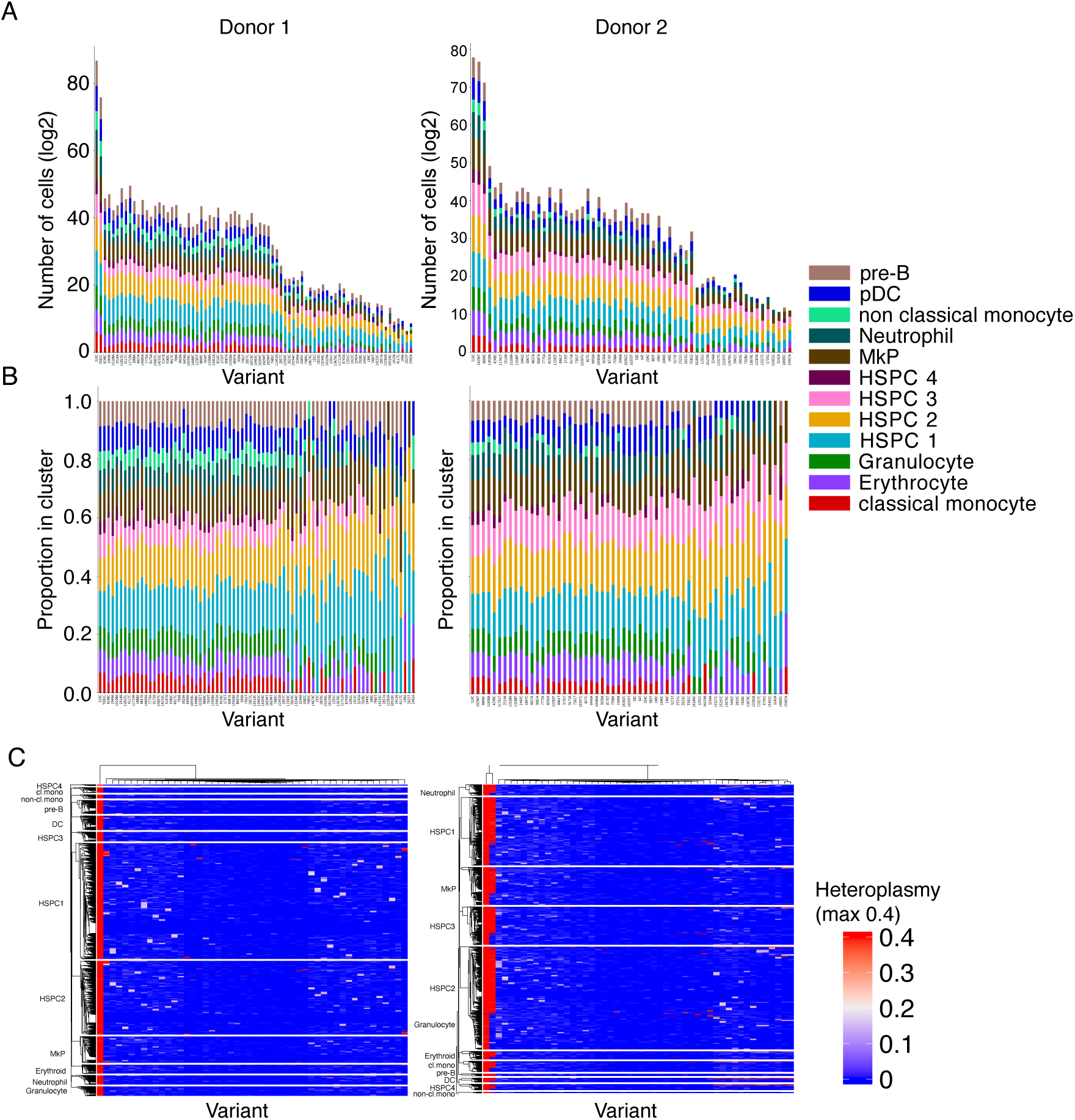
MT barcodes across lineage clusters. (A) Total cell counts (log2) for barcodes (allele frequency>0.01, coverage>10) across hematopoietic clusters. (B) Barcodes (allele frequency>0.01, coverage>10) across hematopoietic clusters normalized within each variant. (C) Cell-by-variant heteroplasmy heatmap for top differentiating variants in Donor 1 and Donor 2, ordered by single-linkage hierarchical clustering within each cell type.

**Figure S7.**
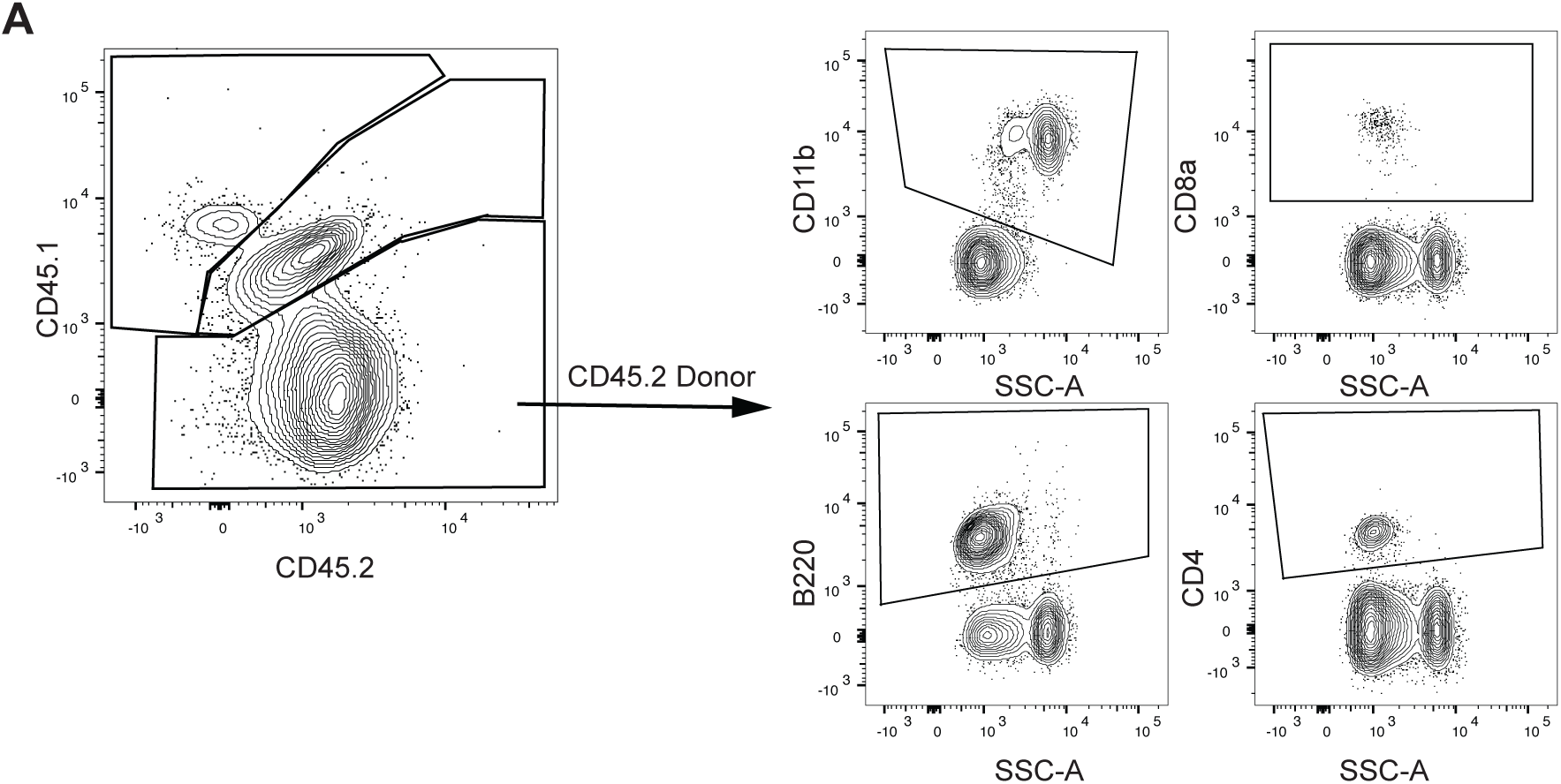
Differentiation of donor, support, and recipient cells in murine transplant experiments. Identification of donor hematopoietic cells (CD45.2) from recipient (CD45.1) and support (CD45.1/CD45.2) cells by flow cytometry to allow for appropriate quantification of donor LT-HSC transplant reconstitution lineage contribution, myeloid (e.g. CD11b), B cell (e.g. B220), T cell (e.g. CD4 and CD8).

**Table S1.** Clone characteristics in CD34+ cells from human donors.

| Donor | Number of clones | Number of cells in clone |  |  | Number of nuclear peaks |  |  | Number of cells in clone (fraction of donor) |  |  |
| --- | --- | --- | --- | --- | --- | --- | --- | --- | --- | --- |
|  |  | mean +/- std | median | max | mean +/- std | median | max | mean +/- std | median | max |
| **Donor 1** | 36 | 101.61 +/- 79.23 | 73 | 429 | 5612.50 +/- 355.74 | 5611 | 6406 | 0.03 +/- 0.02 | 0.02 | 0.12 |
| **Donor 2** | 26 | 100.85 +/- 70.69 | 97.5 | 317 | 5405.59 +/- 531.32 | 5389 | 7457 | 0.04 +/- 0.03 | 0.04 | 0.12 |
| **Donor 3** | 34 | 38.62 +/- 24.36 | 41 | 91 | 3095.06 +/- 644.63 | 3020 | 5184 | 0.03 +/- 0.02 | 0.03 | 0.07 |
| **Donor 4** | 27 | 76.11 +/- 48.80 | 62 | 247 | 3216.47 +/- 366.80 | 3154 | 3892 | 0.04 +/- 0.02 | 0.03 | 0.12 |
| **Donor 5** | 33 | 63.39 +/- 74.85 | 26 | 264 | 3016.17 +/- 517.84 | 3042 | 3895 | 0.03 +/- 0.04 | 0.01 | 0.13 |
| **Donor 6** | 35 | 56.63 +/- 52.50 | 40 | 193 | 3324.59 +/- 492.69 | 3352 | 4498 | 0.03 +/- 0.03 | 0.02 | 0.1 |
| **Donor 7** | 41 | 30.68 +/- 40.60 | 10 | 213 | 3029.99 +/- 567.27 | 2948 | 4437 | 0.02 +/- 0.03 | 0.01 | 0.17 |
| **Donor 8** | 50 | 35.58 +/- 39.73 | 23 | 180 | 2401.89 +/- 573.19 | 2281 | 3902 | 0.02 +/- 0.02 | 0.01 | 0.1 |

## References

1. Pang, W.W., Price, E.A., Sahoo, D., Beerman, I., Maloney, W.J., Rossi, D.J., Schrier, S.L., and Weissman, I.L. (2011). Human bone marrow hematopoietic stem cells are increased in frequency and myeloid-biased with age. Proceedings of the National Academy of Sciences of the United States of America 108, 20012–20017. 10.1073/pnas.1116110108.

2. Sanjuan-Pla, A., Macaulay, I.C., Jensen, C.T., Woll, P.S., Luis, T.C., Mead, A., Moore, S., Carella, C., Matsuoka, S., Bouriez Jones, T., et al. (2013). Platelet-biased stem cells reside at the apex of the haematopoietic stem-cell hierarchy. Nature 502, 232–236.

3. Carrelha, J., Meng, Y., Kettyle, L.M., Luis, T.C., Norfo, R., Alcolea, V., Boukarabila, H., Grasso, F., Gambardella, A., Grover, A., et al. (2018). Hierarchically related lineage-restricted fates of multipotent haematopoietic stem cells. Nature 554, 106–111. 10.1038/nature25455.

4. Grover, A., Sanjuan-Pla, A., Thongjuea, S., Carrelha, J., Giustacchini, A., Gambardella, A., Macaulay, I., Mancini, E., Luis, T.C., Mead, A., et al. (2016). Single-cell RNA sequencing reveals molecular and functional platelet bias of aged haematopoietic stem cells. Nat. Commun. 7, 11075.

5. Pinho, S., Marchand, T., Yang, E., Wei, Q., Nerlov, C., and Frenette, P.S. (2018). Lineage-Biased Hematopoietic Stem Cells Are Regulated by Distinct Niches. Dev. Cell 44, 634–641.e4.

6. Lu, R., Czechowicz, A., Seita, J., Jiang, D., and Weissman, I.L. (2019). Clonal-level lineage commitment pathways of hematopoietic stem cells in vivo. Proc Natl Acad Sci U S A 116, 1447–1456. 10.1073/pnas.1801480116.

7. Busch, K., Klapproth, K., Barile, M., Flossdorf, M., Holland-Letz, T., Schlenner, S.M., Reth, M., Höfer, T., and Rodewald, H.-R. (2015). Fundamental properties of unperturbed haematopoiesis from stem cells in vivo. Nature 518, 542–546. 10.1038/nature14242.

8. Sarrazin, S., Mossadegh-Keller, N., Fukao, T., Aziz, A., Mourcin, F., Vanhille, L., Kelly Modis, L., Kastner, P., Chan, S., Duprez, E., et al. (2009). MafB restricts M-CSF-dependent myeloid commitment divisions of hematopoietic stem cells. Cell 138, 300–313. 10.1016/j.cell.2009.04.057.

9. Muller-Sieburg, C.E., Cho, R.H., Karlsson, L., Huang, J.-F., and Sieburg, H.B. (2004). Myeloid-biased hematopoietic stem cells have extensive self-renewal capacity but generate diminished lymphoid progeny with impaired IL-7 responsiveness. Blood 103, 4111–4118. 10.1182/blood-2003-10-3448.

10. Kirschner, K., Chandra, T., Kiselev, V., Flores-Santa Cruz, D., Macaulay, I.C., Park, H.J., Li, J., Kent, D.G., Kumar, R., Pask, D.C., et al. (2017). Proliferation Drives Aging-Related Functional Decline in a Subpopulation of the Hematopoietic Stem Cell Compartment. Cell Rep 19, 1503–1511. 10.1016/j.celrep.2017.04.074.

11. Hoggatt, J., Mohammad, K.S., Singh, P., and Pelus, L.M. (2013). Prostaglandin E2 enhances long-term repopulation but does not permanently alter inherent stem cell competitiveness. Blood 122, 2997–3000. 10.1182/blood-2013-07-515288.

12. Rundberg Nilsson, A., Soneji, S., Adolfsson, S., Bryder, D., and Pronk, C.J. (2016). Human and Murine Hematopoietic Stem Cell Aging Is Associated with Functional Impairments and Intrinsic Megakaryocytic/Erythroid Bias. PLoS One 11, e0158369. 10.1371/journal.pone.0158369.

13. Frisch, B.J., Hoffman, C.M., Latchney, S.E., LaMere, M.W., Myers, J., Ashton, J., Li, A.J., Saunders, J., Palis, J., Perkins, A.S., et al. (2019). Aged marrow macrophages expand platelet-biased hematopoietic stem cells via Interleukin1B. JCI Insight 5, 124213. 10.1172/jci.insight.124213.

14. Purton, L.E., and Scadden, D.T. (2007). Limiting factors in murine hematopoietic stem cell assays. Cell Stem Cell 1, 263–270. 10.1016/j.stem.2007.08.016.

15. Kiel, M.J., Yilmaz, O.H., Iwashita, T., Yilmaz, O.H., Terhorst, C., and Morrison, S.J. (2005). SLAM family receptors distinguish hematopoietic stem and progenitor cells and reveal endothelial niches for stem cells. Cell 121, 1109–1121.

16. Rector, K., Liu, Y., and Van Zant, G. (2013). Comprehensive hematopoietic stem cell isolation methods. Methods Mol. Biol. 976, 1–15.

17. Balazs, A.B., Fabian, A.J., Esmon, C.T., and Mulligan, R.C. (2006). Endothelial protein C receptor (CD201) explicitly identifies hematopoietic stem cells in murine bone marrow. Blood 107, 2317–2321.

18. Christensen, J.L., and Weissman, I.L. (2001). Flk-2 is a marker in hematopoietic stem cell differentiation: a simple method to isolate long-term stem cells. Proc. Natl. Acad. Sci. U. S. A. 98, 14541–14546.

19. Naik, S.H., Perié, L., Swart, E., Gerlach, C., van Rooij, N., de Boer, R.J., and Schumacher, T.N. (2013). Diverse and heritable lineage imprinting of early haematopoietic progenitors. Nature 496, 229–232. 10.1038/nature12013.

20. Lin, D.S., Tian, L., Tomei, S., Amann-Zalcenstein, D., Baldwin, T.M., Weber, T.S., Schreuder, J., Stonehouse, O.J., Rautela, J., Huntington, N.D., et al. (2021). Single-cell analyses reveal the clonal and molecular aetiology of Flt3L-induced emergency dendritic cell development. Nat Cell Biol 23, 219–231. 10.1038/s41556-021-00636-7.

21. Sun, J., Ramos, A., Chapman, B., Johnnidis, J.B., Le, L., Ho, Y.-J., Klein, A., Hofmann, O., and Camargo, F.D. (2014). Clonal dynamics of native haematopoiesis. Nature 514, 322–327. 10.1038/nature13824.

22. Rodriguez-Fraticelli, A.E., Wolock, S.L., Weinreb, C.S., Panero, R., Patel, S.H., Jankovic, M., Sun, J., Calogero, R.A., Klein, A.M., and Camargo, F.D. (2018). Clonal analysis of lineage fate in native haematopoiesis. Nature 553, 212–216. 10.1038/nature25168.

23. Spencer Chapman, M., Ranzoni, A.M., Myers, B., Williams, N., Coorens, T.H.H., Mitchell, E., Butler, T., Dawson, K.J., Hooks, Y., Moore, L., et al. (2021). Lineage tracing of human development through somatic mutations. Nature 595, 85–90. 10.1038/s41586-021-03548-6.

24. Kim, S., Kim, N., Presson, A.P., Metzger, M.E., Bonifacino, A.C., Sehl, M., Chow, S.A., Crooks, G.M., Dunbar, C.E., An, D.S., et al. (2014). Dynamics of HSPC repopulation in nonhuman primates revealed by a decade-long clonal-tracking study. Cell Stem Cell 14, 473–485.

25. Koelle, S.J., Espinoza, D.A., Wu, C., Xu, J., Lu, R., Li, B., Donahue, R.E., and Dunbar, C.E. (2017). Quantitative stability of hematopoietic stem and progenitor cell clonal output in rhesus macaques receiving transplants. Blood 129, 1448–1457. 10.1182/blood-2016-07-728691.

26. Bowling, S., Sritharan, D., Osorio, F.G., Nguyen, M., Cheung, P., Rodriguez-Fraticelli, A., Patel, S., Yuan, W.-C., Fujiwara, Y., Li, B.E., et al. (2020). An Engineered CRISPR-Cas9 Mouse Line for Simultaneous Readout of Lineage Histories and Gene Expression Profiles in Single Cells. Cell 181, 1410–1422.e27. 10.1016/j.cell.2020.04.048.

27. Scala, S., Basso-Ricci, L., Dionisio, F., Pellin, D., Giannelli, S., Salerio, F.A., Leonardelli, L., Cicalese, M.P., Ferrua, F., Aiuti, A., et al. (2018). Dynamics of genetically engineered hematopoietic stem and progenitor cells after autologous transplantation in humans. Nat. Med. 24, 1683–1690.

28. Lareau, C.A., Ludwig, L.S., Muus, C., Gohil, S.H., Zhao, T., Chiang, Z., Pelka, K., Verboon, J.M., Luo, W., Christian, E., et al. (2021). Massively parallel single-cell mitochondrial DNA genotyping and chromatin profiling. Nat Biotechnol 39, 451–461. 10.1038/s41587-020-0645-6.

29. Bystrykh, L.V., Verovskaya, E., Zwart, E., Broekhuis, M., and de Haan, G. (2012). Counting stem cells: methodological constraints. Nat Methods 9, 567–574. 10.1038/nmeth.2043.

30. Schep, A.N., Wu, B., Buenrostro, J.D., and Greenleaf, W.J. (2017). chromVAR: inferring transcription-factor-associated accessibility from single-cell epigenomic data. Nat. Methods 14, 975–978.

31. van Galen, P., Kreso, A., Wienholds, E., Laurenti, E., Eppert, K., Lechman, E.R., Mbong, N., Hermans, K., Dobson, S., April, C., et al. (2014). Reduced lymphoid lineage priming promotes human hematopoietic stem cell expansion. Cell Stem Cell 14, 94–106. 10.1016/j.stem.2013.11.021.

32. Gekas, C., and Graf, T. (2013). CD41 expression marks myeloid-biased adult hematopoietic stem cells and increases with age. Blood 121, 4463–4472. 10.1182/blood-2012-09-457929.

33. Chapple, R.H., Tseng, Y.-J., Hu, T., Kitano, A., Takeichi, M., Hoegenauer, K.A., and Nakada, D. (2018). Lineage tracing of murine adult hematopoietic stem cells reveals active contribution to steady-state hematopoiesis. Blood Adv 2, 1220–1228. 10.1182/bloodadvances.2018016295.

34. Palchaudhuri, R., Saez, B., Hoggatt, J., Schajnovitz, A., Sykes, D.B., Tate, T.A., Czechowicz, A., Kfoury, Y., Ruchika, F., Rossi, D.J., et al. (2016). Non-genotoxic conditioning for hematopoietic stem cell transplantation using a hematopoietic-cell-specific internalizing immunotoxin. Nat Biotechnol 34, 738–745. 10.1038/nbt.3584.

35. Czechowicz, A., Palchaudhuri, R., Scheck, A., Hu, Y., Hoggatt, J., Saez, B., Pang, W.W., Mansour, M.K., Tate, T.A., Chan, Y.Y., et al. (2019). Selective hematopoietic stem cell ablation using CD117-antibody-drug-conjugates enables safe and effective transplantation with immunity preservation. Nat Commun 10, 617. 10.1038/s41467-018-08201-x.

36. Ansari, A.M., Ahmed, A.K., Matsangos, A.E., Lay, F., Born, L.J., Marti, G., Harmon, J.W., and Sun, Z. (2016). Cellular GFP Toxicity and Immunogenicity: Potential Confounders in in Vivo Cell Tracking Experiments. Stem Cell Rev Rep 12, 553–559. 10.1007/s12015-016-9670-8.

37. Cromer, M.K., Vaidyanathan, S., Ryan, D.E., Curry, B., Lucas, A.B., Camarena, J., Kaushik, M., Hay, S.R., Martin, R.M., Steinfeld, I., et al. (2018). Global Transcriptional Response to CRISPR/Cas9-AAV6-Based Genome Editing in CD34+ Hematopoietic Stem and Progenitor Cells. Mol Ther 26, 2431–2442. 10.1016/j.ymthe.2018.06.002.

38. Ludwig, L.S., Lareau, C.A., Ulirsch, J.C., Christian, E., Muus, C., Li, L.H., Pelka, K., Ge, W., Oren, Y., Brack, A., et al. (2019). Lineage Tracing in Humans Enabled by Mitochondrial Mutations and Single-Cell Genomics. Cell 176, 1325–1339.e22. 10.1016/j.cell.2019.01.022.

39. Miller, T.E., Lareau, C.A., Verga, J.A., DePasquale, E.A.K., Liu, V., Ssozi, D., Sandor, K., Yin, Y., Ludwig, L.S., El Farran, C.A., et al. (2022). Mitochondrial variant enrichment from high-throughput single-cell RNA sequencing resolves clonal populations. Nat. Biotechnol. 40, 1030–1034.

40. Li, H. (2013). Aligning sequence reads, clone sequences and assembly contigs with BWA-MEM. arXiv arXiv:1303.3997.

41. Huang, Y., McCarthy, D.J., and Stegle, O. (2019). Vireo: Bayesian demultiplexing of pooled single-cell RNA-seq data without genotype reference. Genome Biol 20, 273. 10.1186/s13059-019-1865-2.

42. Traag, V.A., Waltman, L., and van Eck, N.J. (2019). From Louvain to Leiden: guaranteeing well-connected communities. Sci Rep 9, 5233. 10.1038/s41598-019-41695-z.

43. Stuart, T., Srivastava, A., Madad, S., Lareau, C.A., and Satija, R. (2021). Single-cell chromatin state analysis with Signac. Nat Methods 18, 1333–1341. 10.1038/s41592-021-01282-5.

44. Waltman, L., and Jan van Eck, N. (2013). A smart local moving algorithm for large-scale modularity-based community detection. The European Physical Journal B 471.

45. Virtanen, P., Gommers, R., Oliphant, T.E., Haberland, M., Reddy, T., Cournapeau, D., Burovski, E., Peterson, P., Weckesser, W., Bright, J., et al. (2020). SciPy 1.0: fundamental algorithms for scientific computing in Python. Nat. Methods 17, 261–272.

